# Endothelial cell polarity and extracellular matrix production rely on functional ATP6AP2 during developmental and pathological angiogenesis

**DOI:** 10.1101/2021.08.16.456486

**Authors:** NR Patel, A Blanks, Y Li, MC Prieto, SM Meadows

## Abstract

The (Pro)renin receptor ((P)RR), also known as ATP6AP2, is a single-transmembrane protein that is implicated in a multitude of biological processes. However, the exact role of ATP6AP2 during blood vessel development remains largely undefined. Here, we use an inducible endothelial cell (EC)-specific *Atp6ap2* knockout mouse model to investigate the role of ATP6AP2 during both physiological and pathological angiogenesis *in vivo*. We observed that postnatal deletion of *Atp6ap2* in ECs results in cell migration defects, loss of tip cell polarity and subsequent impairment of retinal angiogenesis. *In vitro, Atp6ap2* deficient ECs similarly displayed reduced cell migration, impaired sprouting, and defective cell polarity. Transcriptional profiling of ECs isolated from *Atp6ap2* mutant mice further indicated regulatory roles in angiogenesis, cell migration and extracellular matrix composition. Mechanistically, we showed that expression of various extracellular matrix components is controlled by ATP6AP2 via the extracellular-signal-regulated kinase (ERK) pathway. Furthermore, *Atp6ap2* deficient retinas exhibited reduced revascularization in an oxygen induced retinopathy model. Collectively, our results demonstrated a critical role of ATP6AP2 as a regulator of developmental and pathological angiogenesis.

## Introduction

The growth and expansion of the vascular system is a critical physiological process occurring throughout development and adulthood, and pathological conditions can arise with dysregulation of this process. Angiogenesis is the multifaceted process whereby an initial vascular plexus is extended and elaborated throughout the body. A balance in the angiogenic growth of blood vessels is crucial for proper functioning of the vital organs, and both increases and decreases in angiogenesis have been associated with disease states. For example, ischemia secondary to impaired angiogenesis has been implicated in life-threatening disorders such as stroke, myocardial infraction, and neurodegenerative diseases^1–3^. In contrast, excessive angiogenesis is linked to diseases such as diabetic retinopathy and cancer^4,5^. A better understanding of the mechanisms that regulate physiological and pathological angiogenesis will allow us to develop effective therapies that can modulate the angiogenesis process and treat vascular angiogenic-associated disorders.

The murine retina is a well-established model system for studying angiogenesis^6,7^. During retinal angiogenesis, endothelial cells (ECs) at the vascular front rearrange into tip and stalk cells to guide vessel growth that leads to the establishment of the retinal vascular network^8^. The tip ECs are pro-invasive in nature and respond to guidance cues presented in the neighboring tissues and various cell types to facilitate angiogenic sprouting. Tip cell polarity is an important regulator of sprouting angiogenesis, as disruption of EC polarization has been implicated in impaired retinal angiogenesis^9–12^. Furthermore, during the sprouting process, the extracellular matrix (ECM) provides a crucial scaffold for EC attachment and migration^13,14^ and modulates signal-transduction pathways essential for EC morphogenesis^15,16^. Factors that regulate retinal angiogenesis, particularly those involved with EC polarity and ECM deposition are of great interest to the field, though, many of these critical factors remain largely uncharacterized.

The (Pro)renin receptor (P)RR)/ATP6AP2 is encoded by the *Atp6ap2* gene. ATP6AP2 is ubiquitously expressed and highly conserved from mice to humans^17,18^. ATP6AP2 binds renin and prorenin as its natural ligands. The binding of ATP6AP2 to prorenin generates active renin whereas binding to renin amplifies the hydrolysis of angiotensinogen to angiotensin I in the renin angiotensin system (RAS) cascade. These receptor-ligand interactions enhance the activity of the tissue RAS, which is implicated in various pathophysiological conditions^18–21^. However, activation of ATP6AP2 by renin and prorenin also stimulates intra-cellular tyrosine phosphorylation pathways independent of RAS signaling. In addition, ATP6AP2 serves as an adaptor for a number of different proteins, such as the vacuolar-ATPase (v-TPase) and the Wnt receptor complex^22^. These interactions highlight the ligand-dependent and ligand-independent multilevel regulatory properties of ATP6AP2. Accordingly, ATP6AP2 controls a variety of cell biological processes, including autophagy, cell cycle progression, cell polarity, development of kidney vasculature, differentiation of renin-expressing cells, and maintenance of podocytes^23–28^. Deletion of *Atp6ap2* in embryonic stem cells failed to produce viable chimeras upon implantation into blastocysts, confirming the importance of *Atp6ap2* in embryonic viability^29^. Conditional *Atp6ap2* knockout mice were generated to better decipher the roles of ATP6AP2 in normal development and the functioning of different organs^30^. Cell-specific ablations of *Atp6ap2* performed in photoreceptor cells, murine cardiomyocytes, and smooth muscle cells have provided insight into the tissue-specific roles of *Atp6ap2*^25,30,31^. Initial studies reported that *Atp6ap2* regulates angiogenic activity in proliferative diabetic retinopathy^32^. Additionally, *Atp6ap2* was shown to be expressed in infantile hemangioma patients and regulate EC proliferation via the Wnt signaling pathway^33^. *In vitro* studies utilizing human umbilical vein endothelial cells (HUVECs) and implanted matrigel plug assays demonstrated that ATP6AP2 promotes angiogenic properties by inducing the ERK pathway independent of RAS signaling^34^. However, the function of ATP6PA2 in embryonic vascular development and physiological angiogenesis *in vivo* has not been studied and hence remains largely unknown.

Here, we show that *Atp6ap2* is a critical regulator of the angiogenesis process, specifically playing a role in the establishment of the ECM and tip-cell polarity in the retina. Constitutive EC-specific ablation of *Atp6pa2* was embryonic lethal due to vascular defects, while inducible EC-specific deletion of *Atp6ap2* led to defective postnatal retinal angiogenesis. We observed impaired EC polarity in *Atp6pa2* deficient retinas. Moreover, loss of *Atp6pa2* impaired endothelial cell sprouting and migration both *in vivo* and in cultured human ECs *in vitro*. Mechanistically, we established that signaling of ATP6AP2 via the ERK1/2 pathway regulates downstream ECM targets upon binding of prorenin in ECs. Lastly, using an oxygen induced retinopathy (OIR) model, we demonstrated that ATP6AP2 is also critically involved during the revascularization process in pathological angiogenesis.

## Results

### Endothelial-specific deletion of *Atp6ap2* results in impaired angiogenesis *in vivo*

To gain an insight into the role of ATP6AP2 in blood vessel development, we first examined its expression in various tissues by utilizing the available transcriptomic data from several endothelial cell (EC) databases. We used Vascular Endothelial Cell Trans-Omics Resource Database (VECTRDB) to analyze organ-specific, EC expression of *Atp6ap2*^35^. *Atp6ap2* mRNA was detected at similar levels in postnatal day (P)7 brain, liver, lung, kidney, and adult brain ECs, while cultured brain ECs showed the highest levels of *Atp6ap2* expression (Supp. Figure 1a). Examination of single-cell RNA sequencing (scRNA-seq) data generated from P7 isolated brain ECs in the same database showed comparable levels of *Atp6ap2* expression in subtypes of brain ECs, including capillary, venous, arterial and tip cells (Supp. Figure 1b). In addition, we surveyed scRNA-seq data obtained from adult lung and brain ECs and vascular-associated cell types, such as mural cells, perivascular fibroblast-like cells and astrocytes using the following database: http://betsholtzlab.org/VascularSingleCells/database.html^36,37^. Among these data sets, we observed various levels of *Atp6pa2* in the different subtype of lung and brain ECs, as well as their respective vascular support cells (Supp. Figure 1c, d). Evaluation of RNA sequencing (RNA-seq) data from isolated murine retinal ECs at different stages of postnatal development^38^ revealed highest expression levels of *Atp6ap2* at P10 and P15 with retained expression up to P50 (Supp. Figure 1e). Lastly, we performed western blot analysis on different sources of ECs (transformed and primary murine and human ECs) and confirmed that ATP6AP2 is expressed at varying degrees in a multitude of different ECs (Supp. Figure 1f). Overall, these analyses reveal that *Atp6pa2* is expressed in the endothelium, and among various types of ECs, thereby supporting a potential role for *Atp6ap2* in regulating blood vessel growth and maintenance during development and into adulthood, respectively.

To investigate the role of ATP6AP2 during embryonic vascular development, conditional *Atp6ap2*-floxed female mice (*Atp6ap2*^fl/fl^; *Atp6ap2* resides on the X-chromosome) were crossed with male mice expressing a *Cre* transgene under control of the *Tie2* promoter and enhancer^39^ (Supp. Figure 2a). Constitutive EC-specific deletion of *Atp6ap2* resulted in embryonic lethality, as no *Atp6ap2*^fl/Y^;*Tie2-*Cre male offspring were found to be born (Supp. Figure 2b; Y refers to the Y-chromosome). Conversely, *Atp6ap2*^fl/Y^, *Atp6ap2*^fl/wt^ and *Atp6ap2*^fl/wt^;*Tie2-Cre* mice were born and lived throughout adulthood. In order to identify the stages of lethality, timed mating was performed and embryos at multiple stages were analyzed. Gross examination showed that at embryonic day (E) 12.5, *Atp6ap2*^fl/Y^;*Tie2-*Cre embryos exhibited substantial decreases in overall size and an absence of large caliber vessels in the yolk sac and cranial regions as compared to the control genotypes (Supp. Figure 2c). *Atp6ap2*^fl/Y^;*Tie2-*Cre embryos examined at E13.5 and later were in the reabsorption process indicating lethality occurred at approximately E12.5. Therefore, these data suggest that endothelial ATP6AP2 is essential for vascular development. However, lethal vascular defects were less likely to be associated with initial blood vessel formation because major dysfunctions in the vasculogenic process generally result in lethality by E10.5. Instead, the stage of embryonic lethality more closely correlated with defects in angiogenesis, which begins around E9.5 when new vessels form from the initial vascular plexus^40,41^.

To explore the role of endothelial ATP6AP2 during physiological angiogenesis, we generated inducible, EC-specific conditional knockout (iECKO) mice of ATP6AP2 by combining the *Atp6ap2*-floxed and *Cdh5*(PAC)-CreER^T2^ mouse lines^30,42^. The resulting male *Atp6ap2*^fl/Y^;*Cdh5*(PAC)-CreER^T2^ and female *Atp6ap2*^fl/fl^;*Cdh5*(PAC)-CreER^T2^ mice that undergo gene deletion will be further referred to as *Atp6ap2*^iECKO^, while *Atp6ap2*^fl/Y^ and *Atp6ap2*^fl/fl^ mice lacking *Cdh5*(PAC)-CreER^T2^ will be referenced as control mice. Utilizing these mice, our studies centered on the angiogenic blood vessels that form in the retina during postnatal development. Cre-mediated recombination of *Atp6ap2* was induced by daily oral administration of tamoxifen at P1–P3 for early induction, or at P5–P7 for late induction (Figure 1a). Successful, robust EC-specific Cre-mediated deletion of *Atp6ap2* was confirmed at both the mRNA and protein level in isolated lung ECs (iLECs) at P7 (Figure 1b-d). Additionally, the Ai14 (Rosa-CAG-LSL-tdTomato) reporter line^43^ demonstrated EC-specific Cre-mediated recombination in *Atp6ap2*^iECKO^ mice (Supp. Figure 3a, b). Examination of isolectin B4 (IB4)-stained whole mount retinas at P7 following early induction revealed impaired retinal vascular development in *Atp6ap2*^iECKO^ mice. In comparison to control retinas, *Atp6ap2*^iECKO^ retinas exhibited a 17%, 25%, and 43% decrease in vascular outgrowth, density and branchpoints, respectively (Figure 1 e-h). After initial peripheral outgrowth of the superficial vessel plexus, retinal vessels sprout perpendicularly into the deeper retina at approximately P7-8. To assess angiogenesis in the deep layer, delayed inactivation of *Atp6ap2* was performed at P5-P7 and retinas were analyzed at P12. No differences in superficial vascular outgrowth between the control and *Atp6ap2*^iECKO^ retinas were observed (Figure 1j). However, P12 *Atp6ap2*^iECKO^ retinas displayed both reduced vascular density in the superficial vascular plexus (Figure 1k) and severely diminished vascular sprouting into the deep layer of the retina (Figure 1l, m). Taken together, these findings establish that ATP6AP2 is essential for retinal angiogenic growth during postnatal development.

**Figure 1.**
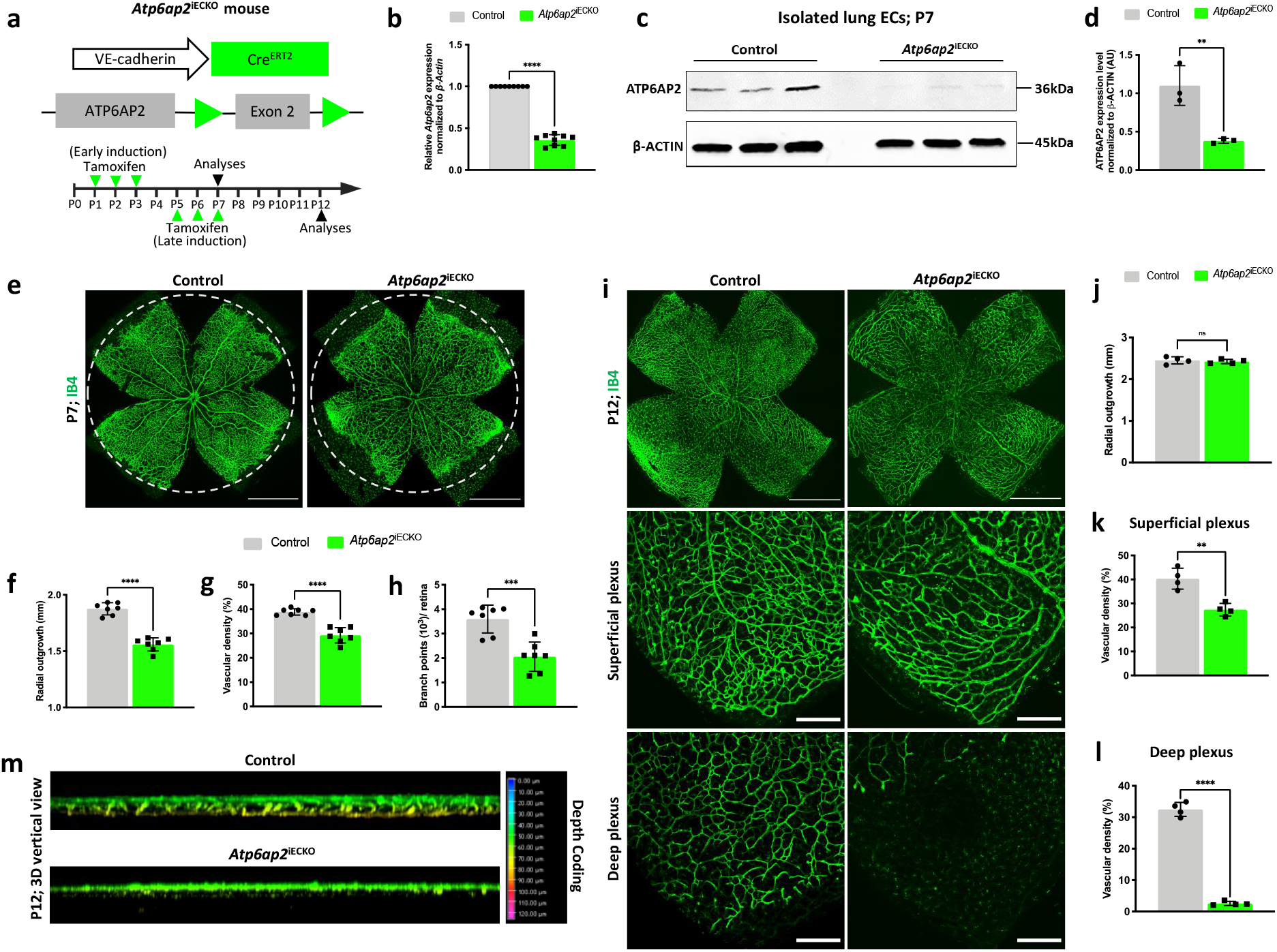
Defective retinal angiogenesis in EC-specific *Atp6ap2* knockout mice. **a**) Strategy for EC-specific deletion of *Atp6ap2* in postnatal mice (*Atp6ap2* induced endothelial cell knockout; *Atp6ap2*^iECKO^) by tamoxifen administration at P1–P3 for early induction or at P5–P7 for late induction. Murine retinas were analyzed at indicated ages. **b**) qPCR analysis of isolated lung ECs (iLECs) show *Atp6ap2* expression levels in control and *Atp6ap2*^iECKO^ mice at P7 (n=3, triplicates for each sample). **c**) **ATP6AP2** Western blot analysis of iLECs from control and *Atp6ap2*^iECKO^ mice. **d**) Densitometric quantification of ATP6AP2 levels in control and *Atp6ap2*^iECKO^ mice at P7 (n=3). **e**) Whole-mount isolectin-IB4 (IB4) stained retinas (dotted circles represent outgrowth in the control retina; scale bars: 1000μm), and quantification of **f**) radial outgrowth, **g**) vascular density and **h**) branch points in control and *Atp6ap2*^iECKO^ mice at P7 (n=7). **i**) Whole-mount IB4 stained retinas (scale bars: 1000μm) and closeup images of IB4^+^ vessels in the superficial plexus and deep plexus (scale bars: 250μm) with quantification of **j**) radial outgrowth (n=4), **k**) vascular density in the superficial plexus (n=4), and **l**) vascular density in the deep plexus (n=4) in control and *Atp6ap2*^iECKO^ mice at P12. **m**) 3D reconstructed images of the IB4^+^ deep vascular plexus and comparisons of the length of the perpendicular growth in control and *Atp6ap2*^iECKO^ mice at P12. Note very few yellow depth-coded vessels in the deep plexus of *Atp6ap2* mutants. Graphs represent mean (bar) ± s.d. (error bars); two-tailed unpaired t-test. ns (not significant; P>0.05), *P<0.05, **P<0.01, ***P<0.001, ****P < 0.0001.

### ATP6AP2 regulates sprouting angiogenesis

The most obvious defects in *Atp6ap2*^iECKO^ retinas were observed at the vascular front where tip cells actively guide angiogenic growth radially. To gain insight into the vascular deficiencies associated with this phenotype, we performed a detailed analysis of the retinal vascular growth front at P7. *Atp6ap2*^iECKO^ mice displayed a decrease in total number of sprouts compared to control retinas (Figure 2a, b). Interestingly, sprouts that did form in *Atp6ap2* mutants showed no difference in filopodia numbers per sprout compared to the control group (Figure 2c, d). To further explore the role of *Atp6ap2* in sprouting angiogenesis, we analyzed sprout formation in a fibrin bead assay using immortalized human aortic endothelial cells (TeloHAECs) treated with siRNAs targeting *Atp6ap2*. First, we verified successful siRNA mediated downregulation at both the mRNA and protein level compared to scrambled control siRNA treated cells (Figure 2e-g). Next, *Atp6ap2* and control siRNA treated TeloHAECs were subjected to the bead sprouting assay. After 120 hours, *Atp6ap2* deficient-ECs displayed defective sprouting with significantly fewer sprouts per bead as compared to control siRNA treated cells (Figure 2h, i). Based on these data, we conclude that ATP6AP2 is necessary for proper sprouting angiogenesis.

**Figure 2.**
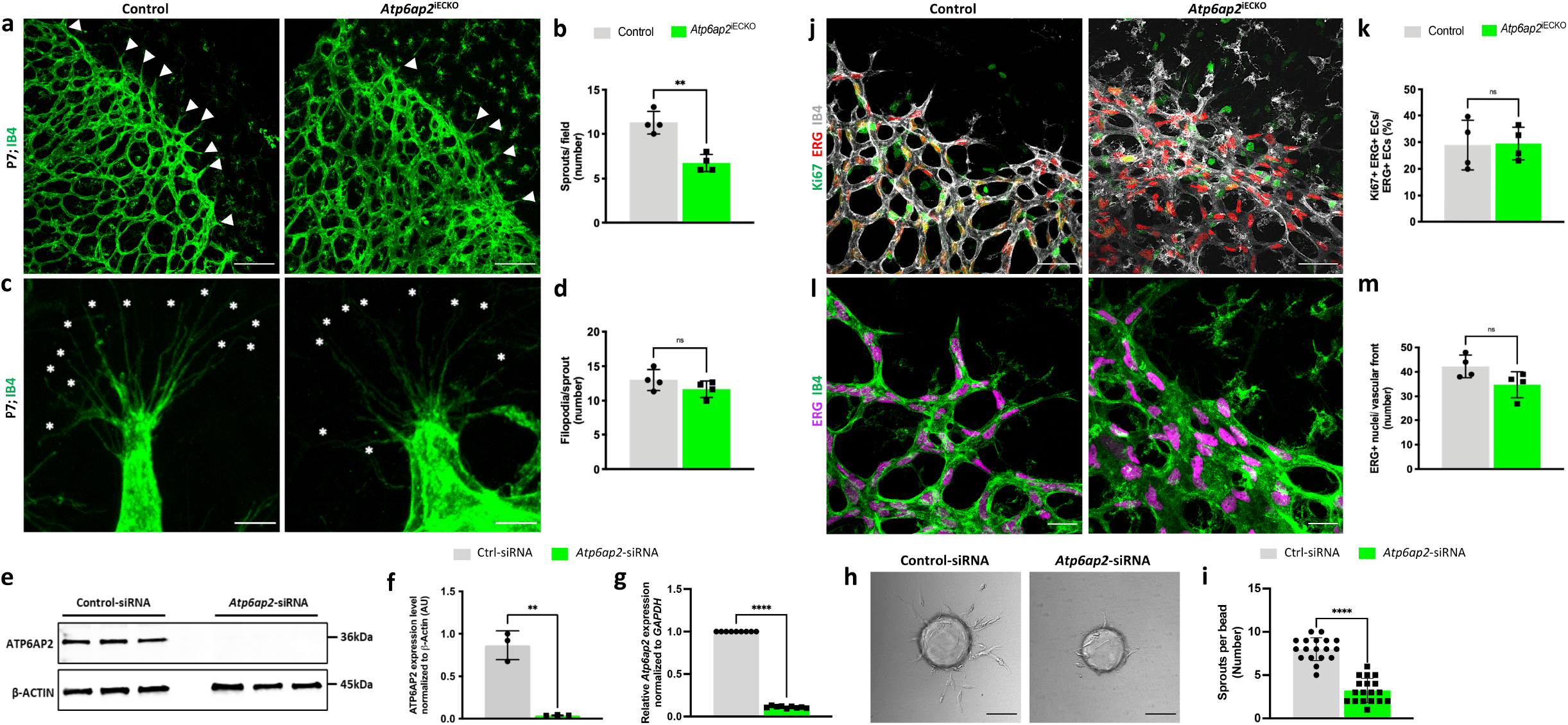
ATP6AP2 is critical for sprouting angiogenesis. **a**) Images of IB4^+^ vessels from P7 control and *Atp6ap2*^iECKO^ retinas with white arrow heads indicating sprouts at the vascular front (scale bars: 100μm). **b**) Quantification of the number of sprouts in control and *Atp6ap2*^iECKO^ mice at P7 (n=4). **c**) Images of IB4^+^ vessels at high magnification showing filopodia in sprouts (white asterisks) at the vascular front (scale bars: 10μm). **d**) Quantification of the number of filopodia per sprout in control and *Atp6ap2*^iECKO^ mice at P7 (n=4). **e**) Western blot analysis of ATP6AP2 and beta-ACTIN (b-ACTIN) in control-siRNA and *Atp6ap2*-siRNA treated TeloHAECs. **f**) Densitometric quantification of ATP6AP2 levels normalized to b-ACTIN in TeloHAECs following siRNA treatments (n=3). **g**) qPCR analysis of control-siRNA and *Atp6ap2*-siRNA treated TeloHAECs for *Atp6ap2* mRNA levels normalized to *Glyceraldehyde-3-Phosphate Dehydrogenase* transcripts (*GAPDH*) (n=3; triplicates for each sample). **h**) Representative images of TeloHAEC sprouting bead-assays embedded in 3D fibrinogen gel at 120 h following control and *Atp6ap2*-siRNA treatments (scale bars: 100μm). **i**) Quantification of the number of sprouts per bead (n=18). **j**) Images of IB4^+^ vessels at the vascular front of control and *Atp6ap2*^iECKO^ P7 retinas immunolabeled for KI67 and the EC-specific nuclear marker ETS transcription factor ERG (scale bars: 50μm). **k**) Quantification of Ki67^+^/ERG^+^ proliferative ECs in control and *Atp6ap2*^iECKO^ mice at P7 (n=4). **l**) Images of ERG^+^ EC-nuclei and IB4^+^ vessels at the vascular front (scale bars: 25μm). **m**) Quantification of the number of ERG^+^ ECs in control and *Atp6ap2*^iECKO^ P7 mice at the vascular front (n=4). Graphs represent mean (bar) ± s.d. (error bars); two-tailed unpaired t-test. ns (not significant; P>0.05), *P<0.05, **P<0.01, ***P<0.001, ****P < 0.0001.

Furthermore, the vasculature in *Atp6ap2* mutants was considerably dense at the growth front, with ECs more closely packed together. Compromised sprouting angiogenesis likely contributed to the increased density, however changes in EC proliferation alone or in combination with impaired sprout formation could also explain this observed phenotype. We assessed whether proliferation in the vascular front was altered by immunofluorescence co-labeling for the ETS-related protein (ERG), an EC-specific nuclear marker, and the proliferation marker KI67. Quantification of KI67^+^/ERG^+^ ECs showed no changes in EC proliferation at the vascular front upon loss of *Atp6ap2* (Figure 2j, k). Moreover, we found no difference in the total number of ERG^+^ ECs at the leading vessel front compared to control Atp6ap2 retinas (Figure 2l, m). Thus, changes in EC proliferation do not appear responsible for the increased vascular density at the growing front.

### Loss of ATP6AP2 leads to defective EC migration and polarity *in vivo* and *in vitro*

In addition to an increased vessel density at the vascular front, *Atp6ap2*^iECKO^ retinas exhibited reduced outgrowth of the superficial vascular plexus (Figure 1e, f). This phenotype often occurs as the result of defective EC migration, which can subsequently cause an increased vessel density at the front. To further evaluate *Atp6ap2’s* role in EC migration, we characterized EC nuclei shape at the vascular front by staining for ERG. We observed that the majority of the nuclei of tip ECs in the control retinas were elliptical, while the nuclei of ECs at the vascular front of *Atp6ap2*^iECKO^ mice were more spherical in shape (Figure 3a, b). Elliptical shaped nuclei are associated with actively migrating cells, whereas spherical shaped nuclei are a characteristic of static cells^12,44,45^. We next performed scratch-wound assays on siRNA treated TeloHAECs to characterize migration *in vitro*. Quantification analysis of the invaded area 40 hours post scratch demonstrated that loss of *Atp6ap2* resulted in impaired wound closure and significantly reduced the rate of EC migration (Figure 3c, d). Together, our *in vivo* and *in vitro* results further support the notion that diminished vascular sprouting and migration in the absence of *Atp6ap2* contribute to reduced retinal vascular outgrowth and increased vessel density at the leading front.

**Figure 3.**
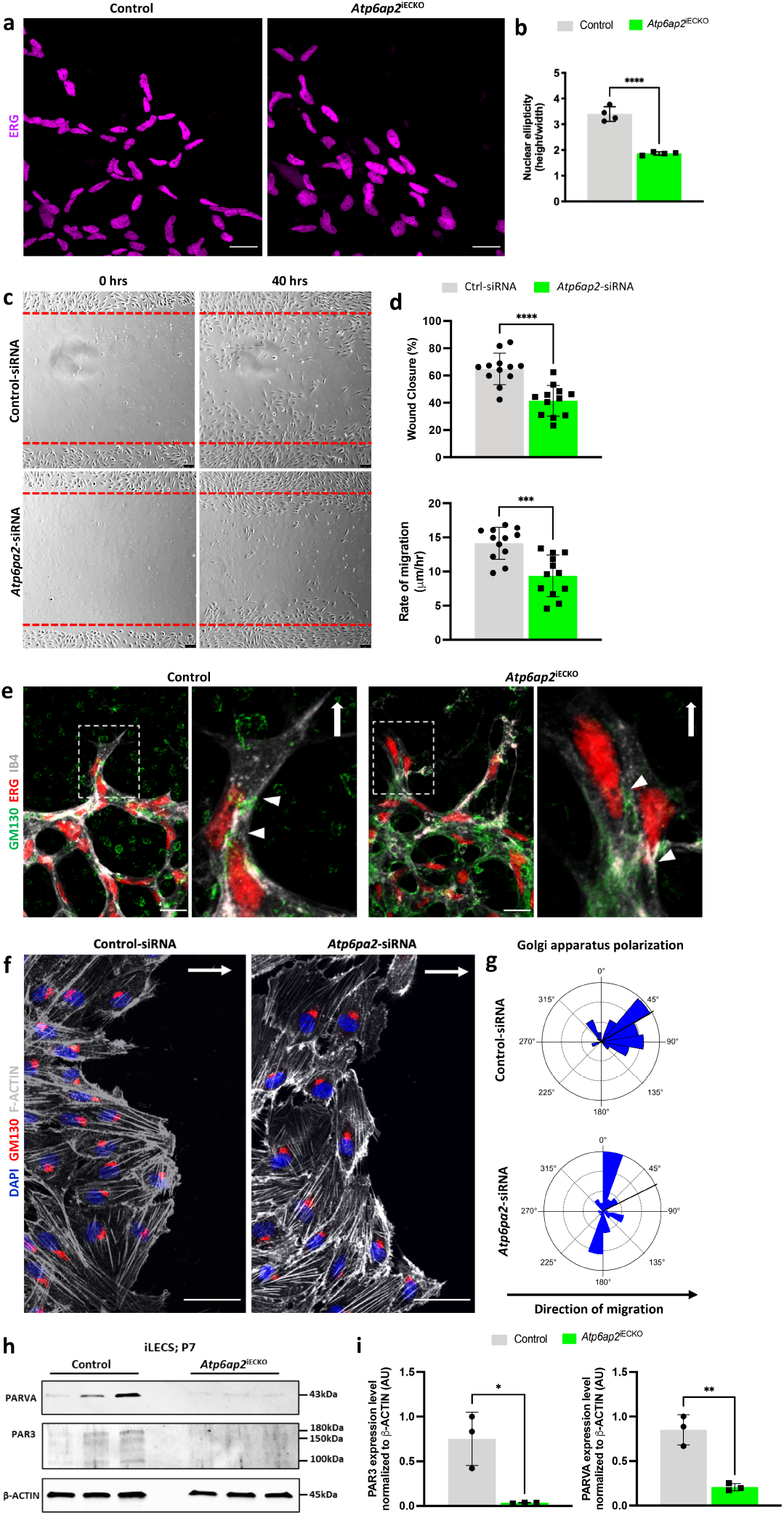
ATP6AP2 regulates endothelial cell migration and polarity *in vivo* and *in vitro*. **a**) Representative images of ERG^+^ nuclei of ECs at the vascular front in P7 control and *Atp6ap2*^iECKO^ retinas (scale bars: 25μm). **b**) Quantification of nuclear ellipticity in control and *Atp6ap2*^iECKO^ mice at P7 (n=4). **c**) Scratch wound assays performed on TeloHAEC monolayers following control and *Atp6ap2*-siRNA treatments. Images at 0 and 40h following the scratch (scale bars: 75μm). **d**) Quantification of the wound closure and rate of migration in control and *Atp6ap2*-siRNA treated TeloHAECs at 40h (n=12). **e**) Images of fluorescently labeled IB4^+^ vessels, ERG^+^ nuclei in ECs and Golgi Matrix Protein GM130^+^ Golgi apparatus at the vascular front with their respective insets highlighting tip cells (white dashed-line boxed) in control and *Atp6ap2*^iECKO^ mice at P7 (scale bars: 25μm). White arrow heads denote the Golgi apparatus position in respect to the nucleus. White arrows indicate direction of migration within the retinal vasculature. **f**) Images of DAPI^+^ nuclei, GM130^+^ Golgi apparatus and F-ACTIN^+^ cytoskeleton of indicated TeloHAECs at 12h after initiating cell migration in a scratch assay (scale bars: 50μm). White arrow indicate the direction of migration. **g**) Rose plots showing the Golgi apparatus polarization in respect to the nucleus in both control and *Atp6ap2*-siRNA treated TeloHAECs at 12h post-scratch (n=52). **h**) Western blot analysis of PAR3, PARVA and b-ACTIN from control and *Atp6ap2*^iECKO^ iLECs at P7 (n=3). **i**) Densitometric quantifications of PAR3 and PARVA levels in control and *Atp6ap2*^iECKO^ iLECs from (**h**). Graphs represent mean (bar) ± s.d. (error bars); two-tailed unpaired t-test. ns (not significant; P>0.05), *P<0.05, **P<0.01, ***P<0.001, ****P < 0.0001.

Defects in directed cell migration can be caused by numerous factors. To further identify potential causes of defective EC migration in *Atp6ap2* mutants, we focused on assessing EC polarity. Alterations in cell polarity have substantial effects on cell migration^10,46,47^, while *Atp6ap2* has been demonstrated to regulate cell polarity dynamics in other cell types^25^. Typically, ECs establish a front-rear polarity axis by orienting their Golgi apparatus in front of the nucleus and towards the direction of migration (towards the periphery). Thus, we initially assessed tip EC polarity at the angiogenic front by staining for IB4, ERG, and Golgi matrix protein 130 (GM130) to define Golgi apparatus polarization in respect to the EC nuclei. In control retinas, tip ECs generally had their Golgi positioned in front of the nucleus towards the peripheral avascular area, while such polarization was rarely observed in the ECs at the growing front of *Atp6ap2*^iECKO^ mice (Figure 3e). This led us to next investigate whether ATP6AP2 regulates endothelial polarization *in vitro*. siRNA treated TeloHAECs were subjected to scratch wound assays and the nucleus-Golgi axis angles were measured near the wound site. The majority of control siRNA treated cells showed polarization of GM130-stained Golgi in front of the nucleus and in the direction of the wound where migration normally occurs (Figure 3f, g). In contrast, *Atp6ap2* siRNA treated cells failed to consistently polarize Golgi towards the wound (Figure 3f, g). These results were consistent with the defective tip-cell polarity observed within *Atp6ap2* deficient ECs *in vivo* and revealed a prominent role for *Atp6ap2* in determining EC polarity.

Previous reports investigating photoreceptor cells have indicated a physical association between ATP6AP2 and PAR3^25^, which has been established to be a critical regulator of polarity in ECs^48,49^ and many other cell types^50,51^. Additionally, ATP6AP2 has been shown to have gene-gene network interactions relating to cancer cell metabolism with PARVA^52^, a member of the Parvins protein family that regulates EC polarity during embryonic blood vessel development^53^. Given these relationships, we examined whether loss of *Atp6ap2* had effects on the *in vivo* expression of either protein. Utilizing proteins from isolated lung ECs (iLECs), we quantified expression levels of PAR3 and PARVA. *Atp6ap2*^iECKO^ mice displayed a substantial reduction in PAR3 and PARVA expression as compared to the control iLECs (Figure 3h, i). Thus, these results connect the *Atp6ap2* deficiency to the downregulation of two major EC polarity determinant proteins. Collectively, our *in vivo* and *in vitro* evidence underscored the importance of ATP6AP2 in the establishment of front-rear EC polarity during angiogenic expansion of the vasculature.

### Gene expression changes in *Atp6ap2* deficient ECs reveal potential roles in various cellular processes

Information pertaining to the function of ATP6AP2 in the endothelium is exceedingly limited. Therefore, we performed RNA-sequencing on iLECs at P7 (Figure 4a) to build an initial, comprehensive understanding of the *Atp6ap2*-associated processes at play in postnatal vascular development. RNA-seq analysis of three biological replicates revealed 1876 differentially expressed genes (823 upregulated genes and 1053 downregulated genes) after *Atp6ap2* inactivation in ECs (Figure 4b). Next, we performed cross-organ comparison of the differentially expressed genes in *Atp6ap2*^iECKO^ mice with those identified as organ-specific EC transcripts^35^. We observed relatively small changes in sets of organ-specific genes for the brain, liver, and kidney upon loss of *Atp6ap2* (Figure 4c). Gene ontology (GO) analysis for enriched biological processes showed upregulated genes to be associated with DNA replication, regulation of cell cycle, and metabolic process (Figure 4d). In contrast, the downregulated genes were enriched in processes associated with angiogenesis, extracellular structure organization, and regulation of cellular response to growth factor stimulus, among others (Figure 4d). Additional heat map analyses indicated downregulation of transcripts encoded by genes involved in promoting angiogenesis (Figure 4e), cell directed migration (Figure 4f) and extracellular matrix (ECM) composition (Figure 4g) in *Atp6ap2*^iECKO^ mice compared to controls. Misregulation in expression of proangiogenic and cell migration genes in *Atp6ap2* mutants further confirmed a vital role of *Atp6ap2* in the regulation of angiogenesis. RNA-seq data also showed downregulation in mRNA expression of the cell polarity-associated gene *α-parvin* in *Atp6ap2* mutants (−0.27 Log_2_fold change). Lastly, GO analyses for phenotype, molecular function, cellular component, and pathway enrichment in differentially expressed genes revealed involvement of downregulated genes with abnormal blood vessel morphology, extracellular matrix structural constituent, cell leading edge, and axon guidance pathway respectively (Supp. Figure 4a-d). Collectively, these transcriptomic findings support our previous data and further establish *Atp6ap2* as having a critical role in multiple processes associated with angiogenesis, including those related to ECM.

**Figure 4.**
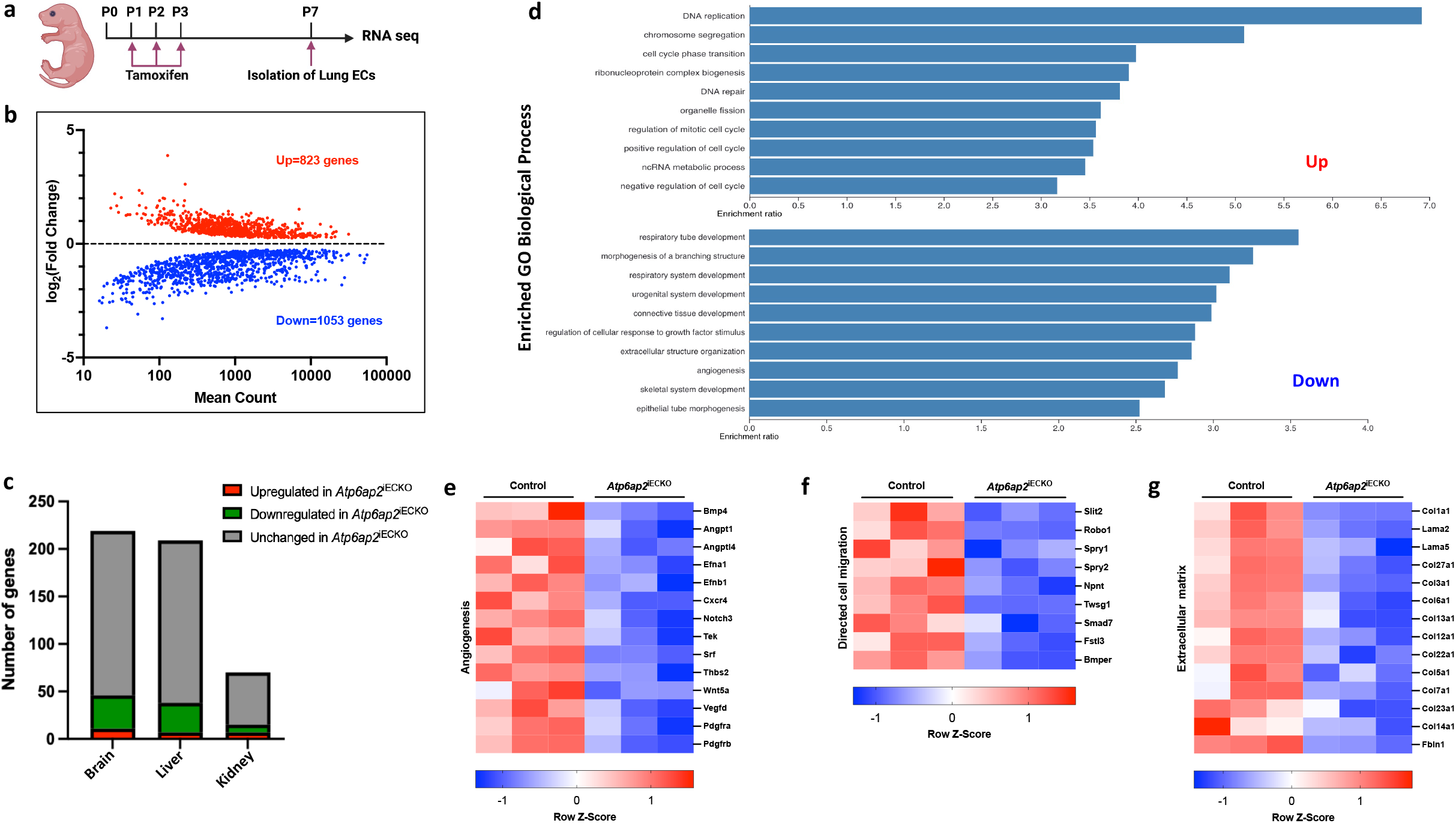
Loss of *Atp6ap2* in ECs results in misregulation of angiogenesis, directed cell migration and extracellular matrix-related genes. **a**) Outline of the workflow used to collect iLECs from control and *Atp6ap2*^iECKO^ mice at P7 (n=3) for RNA-seq analyses. **b**) MA plot of differentially expressed genes between *Atp6ap2*^iECKO^ and control iLECs. Red dots, upregulated genes (823); blue dots, downregulated genes (1053). **c**) Summary of *Atp6ap2*^iECKO^-upregulated and -downregulated EC-gene distribution among organ EC-specific mRNAs. **d**) Top Gene ontology (GO) biological process terms enriched in up- or downregulated genes in *Atp6ap2*^iECKO^ iLECs (False discovery rate (FDR)≤ 0.05). **e-g**) Representative clustered heat maps of gene count Z scores for angiogenesis (**e**), cell migration (**f**), and extracellular matrix-related genes (**g**) that are differentially expressed upon loss of *Atp6ap2* in ECs. Columns represent individual biological replicates.

### ATP6AP2 silencing impairs Prorenin-ERK-mediated pathway and downstream ECM signaling in ECs

A functional ECM is crucial to allow for proper EC adhesion and migration during expansion of the vascular network^54–56^. Our RNA-seq analysis had revealed that ECM-related genes, especially collagens and laminins, were significantly affected by the loss of *Atp6ap2* (Figure 4g). Additionally, we found decreased expression of COLLAGEN III and LAMININ in iLECs of P7 *Atp6ap2*^iECKO^ mice compared to the control (Figure 5 a-c) further validating the RNA-seq results. Together, these data demonstrated that endothelial loss of ATP6AP2 led to a robust reduction in the EC-ECM.

**Figure 5.**
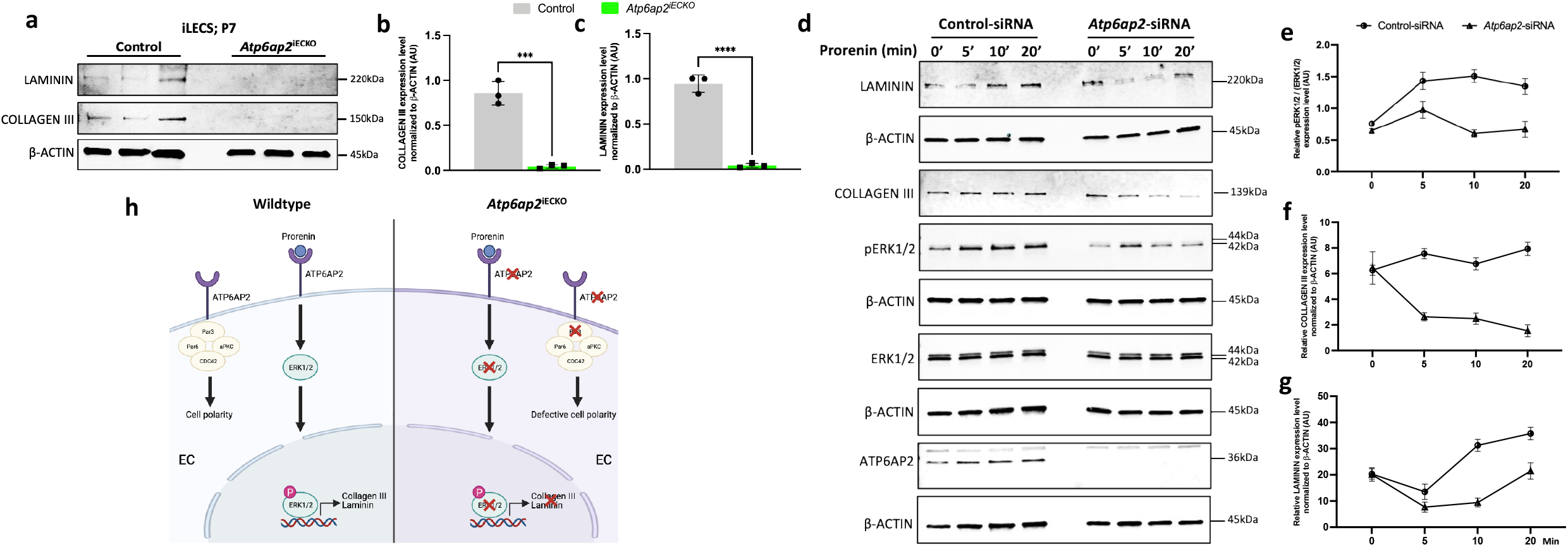
ATP6AP2 knockdown in ECs results in impaired ERK1/2 signaling and downregulation of extracellular matrix proteins. **a**) Western blot analysis of COLLAGEN III, LAMININ and b-ACTIN from control and *Atp6ap2*^iECKO^ iLECs at P7 (n=3). **b, c**) Densitometric quantification of COLLAGENIII (**b**) and LAMININ(**c**) levels in control and *Atp6ap2*^iECKO^ iLECs from (**a**). **d**) Western blot analysis for the indicated proteins in prorenin stimulated TeloHAECs at indicated time points following control and *Atp6pa2*-siRNA treatment. Note that the b-ACTIN blots correspond to the same protein samples represented directly above them. For instance, the b-ACTIN blot at the bottom corresponds to the samples used to assess ATP6AP2. **e-g**) Densitometric quantifications of pERK1/2 (**e**), COLLAGENIII (**f**), LAMININ (**g**) levels in control and *Atp6pa2*-siRNA treated cells at indicated time points following prorenin stimulation (n=3 for each time point). **h**) Working model showing ATP6AP2 regulates extracellular matrix proteins via a mechanism involving ERK1/2 and regulates cell polarity via the PAR3-PAR6 complex in ECs. Graphs represent mean (point/bar) ± s.d. (error bars); two-tailed unpaired t-test. ns (not significant; P>0.05), *P<0.05, **P<0.01, ***P<0.001, ****P < 0.0001.

Prorenin, an inactive precursor of renin, has been shown to bind ATP6PA2 in ECs and subsequently activate the extracellular-signal-regulated kinase (ERK)1/2 pathway independent of the renin-angiotensin system^34^. However, activation of this pathway and its effect on the downstream targets in ECs remain unclear. Interestingly, Prorenin-ATP6AP2 activation of the ERK pathway has been shown to regulate expression of matrix factors in mesangial cells^57^. These findings, coupled with our own results, led us to hypothesize that down regulation of ECM related transcripts observed in *Atp6ap2* deficient ECs is due to impairment in the prorenin mediated activation of ERK in ECs. To address this hypothesis, we stimulated *Atp6ap2*-siRNA treated TeloHAECs with prorenin. Following prorenin stimulation, there were no differences in expression levels of total ERK1/2 in control and *Atp6ap2-*siRNA treated cells (Figure 5d). However, expression levels for phosphorylated ERK1/2 (pERK1/2) were significantly decreased after 5, 10 and 20 minutes of prorenin stimulation in *Atp6ap2*-siRNA treated cells compared to the controls (Figure 5d, e) and as previously reported^34^. Additionally, we examined levels of COLLAGEN III and LAMININ following prorenin stimulation. We observed a consistent reduction of both ECM proteins in *Atp6ap2*-siRNA treated cells compared to the control-siRNA treated cells (Figure 5d, f, g). Based upon these findings, our data point to a mechanism whereby Prorenin-ERK signaling mediates ECM production in the growing endothelium (Figure 5h).

### Loss of *Atp6ap2* results in reduced revascularization in the OIR model

Previous works have shown *Atp6ap2* expression in fibrovascular tissues of murine and human eyes^32,58^ and indicated that ATP6AP2 is linked to patients with proliferative diabetic retinopathy^59^. To expand upon these studies, we examined whether endothelial *Atp6ap2* has a role in pathological angiogenesis in mice by utilizing the oxygen-induced retinopathy (OIR) model, which mimics conditions associated with ocular retinopathies. In these studies, *Atp6ap2* gene deletion was induced at the beginning of the hypoxic phase to assess its role during the neovascularization process (Figure 6a). Analysis of IB4-stained whole mount retinas at P17 showed significantly impaired revascularization in *Atp6ap2*^iECKO^ -OIR mice compared to control-OIR mice (Figure 6b, c). Unlike *Atp6ap2* control-OIR retinas, which displayed small avascular areas and a highly recovered vasculature, *Atp6ap2* mutants exhibited large avascular areas within their retinas. Interestingly, quantification of the neovascular tuft (NVT) areas revealed no differences in tuft formation upon loss of *Atp6ap2* (Figure 6d). In addition, we analyzed EC polarization during the revascularization process in the OIR mice. In control-OIR mice, the Golgi apparatus were generally polarized in front of the nuclei towards the avascular area, similar to P7 control tip cells and control siRNA treated TeloHAECs in the scratch assay (Figure 3e-g). Conversely, Golgi polarization in ECs was disturbed and disorganized in *Atp6ap2*^iECKO^ -OIR mice (Figure 6e), as observed in *Atp6ap2* deficient ECs at the leading edge of the scratch and retina vasculature (Figure 3e-g). These results indicated that ATP6PA2 is essential for establishing EC polarity during the vascular regrowth process in the OIR model.

**Figure 6.**
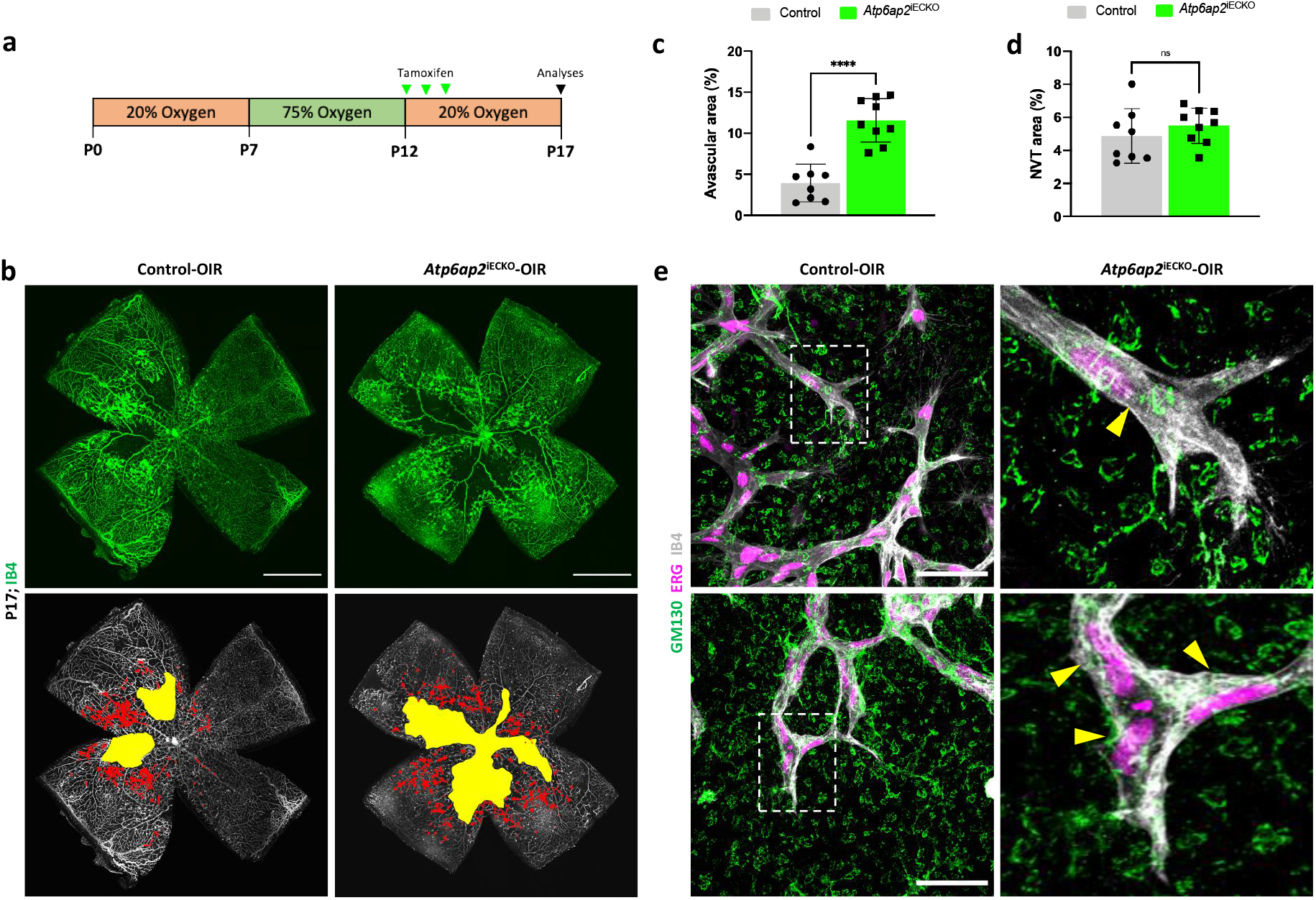
ATP6AP2 is required for proper revascularization during pathological angiogenesis in the OIR mouse model. **a**) Schematic showing the timeline of the OIR protocol with hyperoxia phase from P7-P12, tamoxifen administration from P12-P14, and retina analysis at P17. **b**) Whole-mount IB4 stained retinas showing the avascular area (yellow) and neovascular tuft (NVT) area (red) in control-OIR and *Atp6ap2*^iECKO^-OIR mice at P17 (scale bars: 1000μm). **c**,**d**) Quantification of the avascular area and NVT area in control-OIR (n=8) and *Atp6ap2*^iECKO^-OIR mice (n=9) at P17. **e**) Closeup images of IB4^+^ vessels, ERG^+^ nuclei of ECs, and GM130^+^ Golgi apparatus at the neovascularization edge in control-OIR and *Atp6ap2*^iECKO^-OIR P17 retinas (scale bars; 50μm). The respective insets (white dashed-line boxes) show magnified views of tip cells. Yellow arrow heads indicate the Golgi apparatus position in respect to the nucleus of tip ECs. Graphs represent mean (bar) ± s.d. (error bars); two-tailed unpaired t-test. ns (not significant; P>0.05), *P<0.05, **P<0.01, ***P<0.001, ****P < 0.0001.

## Discussion

ATP6AP2 directs a diverse set of physiological activities in various tissues and cell types. However, the functional role of ATP6AP2 in the endothelium is unclear. In this study, we used endothelial-specific *Atp6ap2* knockout mice to extensively characterize the role of ATP6AP2 in both physiological and pathological angiogenesis during vascular development. Notably, our findings showed that inactivation of *Atp6ap2* in ECs impaired postnatal retinal angiogenesis. Specifically, early deletion (P1-P3) of *Atp6ap2* led to decreased vascular outgrowth in the superficial plexus, while later ablation (P5-P7) disrupted vascularization of the deeper plexus. In addition, transcriptional profiling of *Atp6ap2* deficient ECs displayed a downregulation of genes involved in promoting angiogenesis. Furthermore, the importance of *Atp6ap2* during pathological angiogenesis was demonstrated by the impact that loss of *Atp6ap2* in ECs had on intraretinal vascularization in the OIR model. Together, these results indicate a previously unknown and critical requirement of *Atp6ap2* during *in vivo* angiogenesis and further support an endothelial role for *Atp6ap2* function in pathophysiological settings.

A key finding in this study was the involvement of *Atp6ap2* in regulating tip cell migration and polarity during angiogenic growth. *Atp6ap2*^iECKO^ mice displayed an overall reduced number of sprouts at P7, and the majority of cells at the vascular front displayed spherical-shaped nuclei. These phenotypes are consistent with impaired tip cell migration. Additionally, *Atp6ap2* mutants exhibited defective tip cell polarity. Previously, Hirose et al demonstrated that ATP6AP2 plays a role in neuronal polarization^60^, thus the roles described here for ATP6AP2 in establishing tip EC polarity and regulating polarity-associated gene expression could be shared among different cell types. We observed downregulation of both PAR3 and PARVA in *Atp6ap2*^iECKO^ mice. PAR3 and PARVA are adapter proteins important in establishing and maintaining EC polarity^49,53^. Interactions between ATP6AP2 and PAR3 in retinal homogenates and yeast-two hybrid assays have been reported previously^61^; however, to our knowledge, this is the first report of a potential mechanism involving ATP6AP2 and PARVA in the regulation of EC polarity. A disruption of these suspected interactions in ECs when *Atp6ap2* is lost may be a primary cause of the observed polarity defects (Figure 5h), therefore future studies aimed at understanding this mechanism are warranted. Ultimately, the aberrant EC-polarity observed in *Atp6ap2*^iECKO^ mice is likely responsible, at least in part, for the significantly reduced radial vessel outgrowth, increased vascular density at the growing front, and overall altered vascular architecture. Additionally, it has been shown that in the mouse kidney, conditional deletion of *Atp6ap2* in Foxd1+ stromal cells, which have been indicated as key progenitors in renal arteriole development^62^, is critical for proper formation of the renal artery tree^63^. Thus, ATP6AP2, acting as a functional v-ATPase subunit, contributes to the morphogenetic regulation of the renal arterial network in a non-EC autonomous manner. Therefore, additional roles of ATP6AP2 in various cell types that affect blood vessel formation should be taken into consideration as well. Moreover, considering the many reported functions of ATP6AP2, additional processes, such as actin cytoskeletal and microtubule dynamics, energy metabolism, or vesicular acidification, could also factor into these phenotypes and remain potential areas for further investigation.

Our studies also uncovered a connection between ATP6AP2 and the ECM of the endothelium. RNA-seq analyses revealed downregulation of ECM genes in response to loss of *Atp6ap2* in ECs. Mechanistically, we showed that decreases in ECM protein levels, specifically Collagen type III and Laminin, were regulated by ATP6AP2 via the ERK1/2 pathway. Collagen and Laminin are each implicated in the regulation of different phases in the angiogenesis process. For instance, exposure of microvascular ECs to Collagen-1 leads to the formation of precapillary cords, whereas these same cells do not undergo a morphology change when exposed to Laminin-1^16,64^. Furthermore, it is proposed that collagen is important for EC invasion and lumen formation during angiogenesis, while Laminin is crucial during the vessel maturation and stabilization phase. The direct effect of ECM proteins on EC function during angiogenesis, neovascularization, and blood vessel maturation highlights the need for investigation into their regulation. In fact, we expect that loss of ECM in the endothelium contributes to the various phenotypes observed in *Atp6ap2* mutant mice, but the mechanism and extent of their contribution are currently unknown. Overall, changes in the ECM and EC polarity in our model were not entirely surprising, as ATP6AP2 has been implicated in ECM and cell polarity regulation in multiple cell types^61,65^. Accordingly, our findings suggested that ATP6AP2 may possess similar and/or conserved roles among various cell types, which could provide functional context for studying different biological processes and diseases.

Although not fully investigated in these studies, our RNA-seq data revealed a number of critical angiogenic genes that were significantly downregulated in *Atp6ap2* deficient ECs (Figure 4e). For example, the vascular ligand angiopoietin-1 (*Angpt1*) and its corresponding receptor tyrosine kinase Tek (also known as Tie2) are crucial for multiple vascular processes, including sprouting angiogenesis and EC migration^66^, which were investigated in this study. Moreover, Angpt1 and Tek are important in EC permeability, quiescence and maturation^66^. From a pathological standpoint, mutations in Tek lead to venous malformations in humans^67^, and angiopoietin-Tek signaling has a significant role in tumor angiogenesis.^66^ As another example, Notch3, one of four Notch receptors involved in the Notch signaling pathway, is important for vascular smooth muscle cell (VSMC) development and function. Postnatal Notch3 null mutants form structurally defective smooth muscle cell layers around arterial vessels due to deficiencies in VSMC maturation and arterial differentiation^68^. Furthermore, adult Notch3^-/-^ mice exhibit VSMC degeneration in the retina and brain causing hemorrhage and loss of vessel integrity and blood brain barrier function^69^. Interestingly, platelet-derived growth factor receptor-beta (PDGFRb), another transcript significantly decreased in our *Atp6ap2* mutant ECs (Figure 4e), is intimately involved in VSMC biology and its expression is regulated by Notch3 signaling^70^. Notch3 also regulates vascular tone and flow-mediated dilation of arterial vessels, and adult Notch3 mutant mice show an increased incidence of developing heart failure under angiotensin II-induced hypertensive conditions^69^. Therefore, based on the findings in our transcriptional profiling studies, it can be speculated that endothelial *Atp6ap2* plays prominent roles in major health-related vascular processes, such as vessel regeneration/remodeling, tumor angiogenesis, vascular malformation, hypertension, and diabetes mellitus. Understanding the cellular and molecular mechanisms involved in the role of ATP6AP2 in angiogenesis is the first step to advance these medically relevant fields.

Prorenin, a ligand for ATP6AP2, is elevated in the plasma of patients with microvascular complications related to diabetic retinopathy^32,71^. Although systemic prorenin levels, are typically lower than the pharmacological threshold for (P)RR activation, this scenario might differ during high glucose conditions^72^. In example, during high glucose conditions, as seen in diabetic patients, there are trafficking alterations of ATP6AP2, further increasing the physical interaction between (P)RR and prorenin, which leads to upregulation of downstream fibrotic factors^73^. This process is clinically relevant due to the potential therapeutic effects of targeting this receptor with an inhibitor. Blocking ATP6AP2 activity prevented cardiac fibrosis and diabetic retinopathy in hypertensive and diabetic rats, respectively^21,74,75^. However, inhibition of ATP6AP2 during pregnancy and/or later on in life may not be an ideal treatment option clinically, as its roles in many physiological processes are not completely understood. Our study elucidated the potential role of ATP6AP2 in vascular angiogenesis and showed that loss of *Atp6ap2* had an overall detrimental effect on the growth, organization, and function of the endothelium with serious pathological events. Therefore, systemic inhibition of ATP6AP2 may have unanticipated consequences since blood vessels are pervasive in nearly all tissues. Future research targeting ATP6AP2 for the treatment of *Atp6ap2-* related disorders will need to consider disease settings and tissue specificity of ATP6AP2.

In conclusion, this study defines the role of endothelial *Atp6pa2* during postnatal retinal angiogenesis and its involvement during neovascularization in the OIR model. Importantly, we identified ATP6AP2 as a regulator of EC polarity during physiological and pathological angiogenesis. Additionally, we found that loss of *Atp6pa2* leads to downregulation of ECM proteins due to impaired signaling in the ERK1/2 pathway in ECs. These findings provide the first evidence to our knowledge, of an association between endothelial cell-type specific *Atp6ap2* and the morphogenetic, well-characterized angiogenesis process that establishes the retinal vascular network.

## Materials and Methods

### Mice and treatment

All animal experiments were performed in accordance with Tulane University’s Institutional Animal Care and Use Committee policy. Floxed *Atp6ap2* (*Atp6ap2*^fl/Y^) mice were crossed with two endothelial-specific Cre lines: *Tie2-cre* for embryonic and *Cdh5-Cre*^*ERT2*^ for postnatal studies, respectively^39,42^. For embryonic studies, timed mating was carried out, designating embryonic day 0.5 (E0.5) as noon on the day a vaginal plug was observed. To induce Cre recombination postnatally, newborn offspring were administered tamoxifen (Sigma, T5648) orally at a concentration of 100 μg on postnatal days 1 to 3 (P1–P3) or 5 to 7 (P5–P7) and the retinas analyzed at P7 or P12, respectively.

### Immunofluorescence analysis of mouse retina

Whole mount staining of retinas was performed as previously described^76^. Briefly, eyeballs were removed and fixed in 4% PFA in PBS for 1 h at 4°C. Eyeballs were washed three times in 1X PBS, retinas were dissected, permeabilized in 1% TritonX-100/PBS (PBST) at room temperature for 30 min, and blocked with CAS-block (Thermofisher, 88120) at room temperature for 30 min. Primary antibodies were diluted in CAS-block and incubated at 4°C overnight, followed by incubation in appropriate secondary antibodies for 4 h at room temperature. Retinas were washed in three times in 1X PBS and mounted on a slide using the ProLong™ Diamond Antifade Mountant (Fisher scientific, P36961). Antibodies: IB4 (Thermofisher, I21411, 1:250), Anti-RFP (Abcam, ab62341), ERG1/2/3 (Abcam, ab196149, 1:200), GM130 (BD, 610822, 1:100), and KI67 (Cell signaling, 9449, 1:100).

### Morphological analysis of retinal vasculature

Retinal vasculature analysis was performed using ImageJ and Angiotool analysis software^77,78^. Vascular outgrowth was measured as the distance from the optical nerve to the periphery in each leaflet of the retina and averaged. Vascular density and the number of branching points were quantified using whole retinal images in the Angiotool analysis software. The number of sprouts and filopodia were counted manually at the vascular front in 20x and 100x images respectively. Quantification of ERG^+^ nuclei of ECs was performed at the vascular front of 60x image using ImageJ. Nuclear ellipticity was determined by dividing the measured nuclear height by nuclear width of 10-12 ERG^+^ cells per image at the vascular front and presented as a ratio. For cell proliferation analysis, ERG^+^/KI67^+^ cells were quantified at the vascular front of 40x image using ImageJ.

### Isolation of murine lung endothelial cells

Isolation of murine lung ECs was performed as previously described^79^. Briefly, lungs were harvested at P7, minced, and digested in Collagenase-I (Fisher scientific, 17-100-17)/Dispase (Corning, 354235) buffer at 37°C for 30 min under constant rotation. The digested tissue was triturated ten times through 18G needle to dissociate the clumps and filtered using a 70μm nylon mesh (VWR, 10199-656) to remove any cell debris. The cell suspension was centrifuged at 1200 rpm, 4°C for 5 min. Single cell suspension were incubated with sheep anti-rat IgG dynabeads (Invitrogen, 11035) coated with CD31 antibody (BD Pharmingen, 553370) at room temperature for 20 min with agitation. Lastly, CD31+ cells were isolated using a magnetic rack separator and either RNA or protein was isolated from the cells for downstream analysis.

### Western blot

Protein was isolated from the cells using the RIPA lysis buffer (Thermo, 89901) supplemented with protease inhibitor cocktail (Thermofisher, 78430) and phosSTOP (Sigma, 04906845001). Cell lysates were centrifuged at 13,000 X *g* for 10 min at 4°C and supernatants were collected. Protein concentrations were determined using the Pierce™ Coomassie (Bradford) protein assay kit (Thermo, 23200). Laemmli buffer (Biorad, 1610737) supplemented with β-mecrapto-ethanol (Biorad, 1610710) was added to the protein samples and boiled at 95°C for 5 min for denaturation. Each protein lysate was separated on 4-20% Mini-PROTEAN TGX Precast gels (Biorad, 4568094) and blotted onto 0.2 µm PVDF membrane (Biorad, 1704156). Membranes were blocked for 30 min with 5% BSA in TBST (0.1% Tween 20 in TBS) and incubated in primary antibodies diluted in the blocking buffer at 4°C overnight. Following primary antibody incubation, membranes were washed in TBST and incubated with appropriate secondary antibodies for 1h at room temperature. Target proteins were detected using the LI-COR Odyssey imaging system. Band densitometry was quantified using the ImageJ software.

The following primary antibodies were used for the western blot analysis: Anti-β-Actin (Cell signaling, 3700S, 1:5000), anti-Atp6ap2 (Sigma, HPA003156, 1:1000), anti-p44/42 MAPK (Erk1/2) (Cell signaling, 4695, 1:1000), anti-phospho-p44/42 MAPK (pErk1/2) (Cell signaling, 9101, 1:1000), anti-Collagen type III (Abcam, ab184993, 1:1000), anti-Laminin (Sigma, L9393, 1:1000), anti-Partitioning-defective 3 (Par3) (Sigma, 07-330, 1:1000), anti-Parvin (Cell signaling, 8190, 1:1000).

### Quantitative polymerase chain reaction

Total RNA was extracted from the samples using the GeneJET RNA Purification Kit (Thermofisher, K0732) following the manufacturer instructions. To determine the mRNA expression levels, 1 μg of extracted RNA was transcribed into cDNA using the iScript Reverse Transcription Supermix kit (Biorad, 1708840). Quantitative RT-PCR was performed using the PerfeCTa SYBR Green SuperMix (Quantabio, 95071) on CFX96 system (Biorad) following manufacturer instructions. The list of qRT–PCR primers used in this study is included in Supplemental Table 1. Relative gene expression was determined using the ΔΔCt method. Three independent biological replicates were used, and three technical replicates were performed per sample.

### Cell culture and stimulation

Telomerase human aortic endothelial cells (TeloHAECs) (ATCC, CRL-4052) were cultured in EBM-2 basal media (Lonza, 3156) supplemented with EGM-2 Bullet kit (Lonza, 3162) at 37°C and 5% CO_2_. For stimulation studies, cells transfected with *Atp6ap2-*siRNA or control-siRNA were serum starved overnight and stimulated with 20nM prorenin (Cayman chemical, 10007599) for indicated time points. Following stimulation, protein was extracted from each treatment for analysis via immunoblotting.

### RNA interference

TeloHAECs were transfected with a pool of *Atp6ap2* (Dharmacon, L-013647-01-0005) and non-targeting control-siRNAs (Dharmacon, D-001810-10-05) using Lipofectamine 3000 (ThermoFisher, L3000015) following manufacturer instructions. The final concentration of siRNA solution was 200nM. Cells were harvested and analyzed for knockdown efficiency 48h after transfection.

### Scratch wound healing assay

Scratch assay was performed on confluent control and *Atp6ap2-*siRNA transfected TeloHAECs. A horizontal and vertical scratch was created in each well using a 200μl pipet tip followed by two washes with EGM-2 media. Phase contrast images were taken right after the scratch (0 h) and 40 h after. Percentage of wound closure and rate of migration was analyzed using the ImageJ software.

### Cell staining and polarity analyses

TeloHAECs were cultured on gelatin-coated coverslips. Cells were stained 12 h following the scratch for polarity studies. For staining, cells were fixed in 4% PFA for 10 min, permeabilized in 0.1% NP40/PBS for 15 min, and blocked in CAS-block for 30 min at room temperature. Cell-coated coverslips were incubated in primary antibodies diluted in CAS-block for 1 h at room temperature, secondary antibodies and DAPI diluted in CAS-block for 1 h at room temperature. After several washes in 1X PBS, coverslips were air-dried and mounted on a slide using the ProLong™ Diamond Antifade Mountant. After mounting, random fields of view for each treatment were imaged with a Nikon confocal microscope (40x objective). Cell polarity towards the scratch was determined by measuring nucleus-Golgi apparatus axis angle in 10-12 cells per image from three independent experiments using ImageJ software. Antibodies used for cell immunofluorescence analysis were: DAPI (Invitrogen, R37606), Phalloidin (F-Actin) (Invitrogen, A12379, 1:200), and GM130 (Cell signaling, 12480, 1:100).

### 3D fibrin gel bead sprouting assay

The 3D in vitro bead assay was performed and analyzed as previously described^80^. Briefly, siRNA transfected TeloHAECs were coated onto Cytodex 3 beads (Cytivia, 17048501) at a concentration of 400 cells per bead in 1ml of EGM2 media incubated for 4 h at 37 °C with gentle shaking every 20 min, transferred to a 25cm^2^ flask, and incubated overnight in 7 ml of EGM2 media at 37°C and 5% CO_2_. Next day, TeloHAEC-coated beads were resuspended in 2.5mg/ml fibrinogen (Millipore, F8630) in PBS. Aprotinin (Millipore, A1153) was added to the fibrinogen/bead solution at a concentration of 4U/ml. Thrombin (Millipore, 605157) was added to the center of a well at a concentration of 50U/ml followed by the fibrinogen/bead solution in a glass-bottom 24-well plate. The gels were allowed to solidify for 5 min at room temperature before it was placed at 37 °C and 5% CO_2_ for 30 min. Normal human lung fibroblasts (Lonza, 2512) were plated on top of the fibrin gel at a concentration of 10,000 cells per well. Media was changed every other day and sprouts were analyzed 3-5 days following sprout formation.

### RNA sequencing and gene expression analysis

Total RNA was extracted from isolated lung ECs and quantified using Qubit RNA High Sensitivity Assay Kit (Thermofisher, Q32852). RNA integrity was determined using the Bioanalyzer RNA 6000 Nano assay kit (Agilent, 5067-1511). RNA library construction was performed with the TruSeq RNA Library Prep Kit v2 (Illumina, RS-122-2001) according to the manufacturer instructions. The resulting mRNA library was quantified using Qubit dsDNA High Sensitivity Assay Kit (Thermofisher, Q32851) and verified using the Bioanalyzer DNA1000 assay kit (Agilent, 5067-1505). Verified samples were sequenced using the NextSeq 500/550 High Output Kit v2.5 (150 Cycles) (Illumina, 20024907) on a Nextseq 550 system (Illumina, SY-415-1002). Sequenced reads were aligned to the mouse (mm10) reference genome with RNA-seq alignment tool (version 2.0.1). The aligned reads were used to quantify mRNA expression and determine differentially expressed genes using the RNA-seq Differential Expression tool (version 1.0.1). Both alignment and differential expression analysis were performed using the tools in the BaseSpace Sequence Hub. Over-representation analysis of differentially expressed genes was performed using the WEB-based Gene SeT AnaLysis Toolkit (WebGestalt)^81^. Sequencing data have been deposited in the Gene Expression Omnibus (GEO) database with accession code (GSE179431).

### OIR studies

To induce retinal pathological angiogenesis, the OIR model was used, as previously described^82^. From P7 to P12, pups were placed in hyperoxia chamber (Biospherix, ProOx110) with 75% oxygen; at P12, pups were returned to room air until P17. Tamoxifen was administered to pups at a concentration of 200μg from P12–P14 to induce Cre recombination. Retinas were collected at P17 and processed as outlined in the section of immunofluorescence analysis of mouse retina. Quantification of the avascular and neovascular tuft (NVT) area was performed using an automated OIR retinal image analysis software^83^.

### Statistical analysis

Data analysis was performed using GraphPad Prism version 9.0.0 for MacOS, GraphPad Software, San Diego, California USA, www.graphpad.com. Quantified data are presented as bar graphs of mean± standard deviation., shown as error bars or as rose diagrams of the mean. Unpaired two-tailed Student’s t tests assuming equal variance were used to determine statistical significance between two groups with a *p*-value < 0.05 was considered statistically significant.

## Author Contributions

N.R.P. and S.M.M developed and designed the experiments. N.R.P, A.B and Y.L. performed the experiments. N.R.P., M.C.P and S.M.M wrote the manuscript.

## Acknowledgements

This work was supported by Tulane University start-up funds (S.M.M), the Department of Defense (PRMRP-160198) (S.M.M), and the NIH-DK104375 grant (M.C.P). We would like to thank Jovanny Zabaleta and Jone Garai at The Louisiana Cancer Research Consortium (LCRC) Translational Genomics Core Center for their continued support with our RNA-seq experiments.

