## Supplemental files for "Endothelial cell polarity and extracellular matrix production rely on functional ATP6AP2 during developmental and pathological angiogenesis"

Supplementary Figure legends:

**Supplementary Figure 1. ATP6AP2 is robustly expressed in vascular endothelium. a**) Analysis of *Atp6ap2* expression levels in non-EC, ECs and total tissue of P7 brain, kidney, liver, lung, and adult brain using the Vascular Endothelial Cell Trans-omics Resource Database (VECTRDB). **b**) Analysis of *Atp6ap2* expression levels in P7 isolated brain ECs and its subtypes using single cell RNA-seq data in VECTRDB. **c,d**) *Atp6ap2* expression levels in adult murine brain and its support cells (**c**) and lung and its support cells (**d**), as determined using the single cell RNA-seq data from the following database: http://betsholtzlab.org/VascularSingleCells/database.html. Brain data: PC - Pericytes; SMC - Smooth muscle cells; MG - Microglia; FB - Vascular fibroblast-like cells; OL - Oligodendrocytes; EC - Endothelial cells; AC - Astrocytes; v - venous; capil- capillary; a - arterial; aa - arteriolar; 1,2,3- subtypes. Lung data: FB - Vascular fibroblast-like cells; CP - Cartilage perichondrium; PC - Pericytes; VSMC - Vascular smooth muscle cells; EC - Endothelial cells; capil - capillary; a - arterial; c - continuum; L - Lymphatic; 1,2,3,4 – subtypes. **e**) Expression levels of *Atp6ap2* in murine retinal endothelial cells during postnatal development determined using the available bulk RNA sequencing data[35]. **f**) Western blot analysis of ATP6AP2 and b-ACTIN expression in various murine and human primary and transformed ECs.

**Supplementary Figure 2. Endothelial-specific deletion of *Atp6pa2* induces embryonic lethality. a**) Strategy for EC-specific deletion of *Atp6ap2* during embryogenesis using the *Tie2*-Cre driver line and summary of the timepoint when vascular defects were observed. **b**) Genotype frequencies of postnatal day 21 pups from *Tie2*-Cre and *Atp6ap2* $X^f/X^f$ mating pairs. No postnatal deaths were observed in pups that were born. **c**) Whole mount, bright-field images of *Atp6ap2* $X^f$/Y and *Atp6ap2* $X^f$/Y;Tie2-cre yolk sacs and embryos at E12.5. Note the absence of large perfused vessels within the yolk sac and brain and reduced size of *Atp6ap2* $X^f$/Y;Tie2-cre embryos (scale bars: 2mm).

**Supplementary Figure 3. Inducible EC-specific Cre recombination in *Atp6ap2*$^{iECKO}$ mice. a**) Schematic representation of the experimental timeline for monitoring EC-specific Cre recombination using the Rosa-tdTomato reporter integrated in *Atp6ap2*$^{iECKO}$ mice, which harbor VE-cadherin Cre-ERT2. **b**) Whole-mount images of control and *Atp6ap2*$^{iECKO}$ retinas showing $IB4^+$ vessels and tdTomato expression only in the ECs of *Atp6ap2*$^{iECKO}$ mice (scale bars: 1000μm).

**Supplementary Figure 4. Gene ontology (GO) analysis of differentially expressed genes following loss of *Atp6ap2* in ECs. a-d**) Top terms enriched in upregulated and downregulated genes associated with (**a**) phenotype, (**b**) molecular function, (**c**) cellular component, and (**d**) pathway following RNA-seq analysis of *Atp6ap2*-deficient iLECs.

**Supplementary Table 1 List of qPCR primer sequences**

### Supplementary figure 1

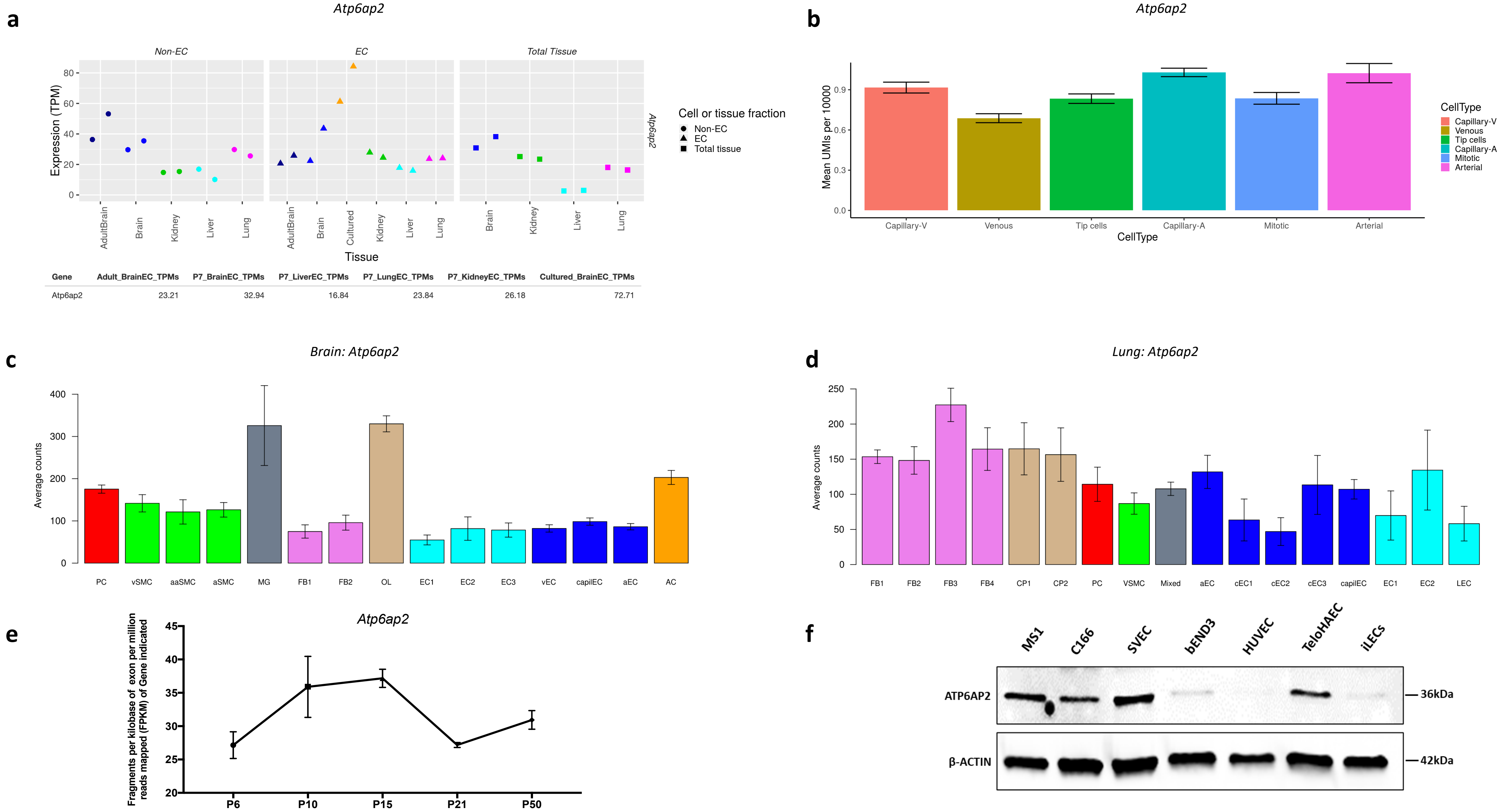


| Gene | Adult_BrainEC_TPMs | P7_BrainEC_TPMs | P7_LiverEC_TPMs | P7_LungEC_TPMs | P7_KidneyEC_TPMs | Cultured_BrainEC_TPMs |
|---|---|---|---|---|---|---|
| Atp6ap2 | 23.21 | 32.94 | 16.84 | 23.84 | 26.18 | 72.71 |

### Supplementary figure 2

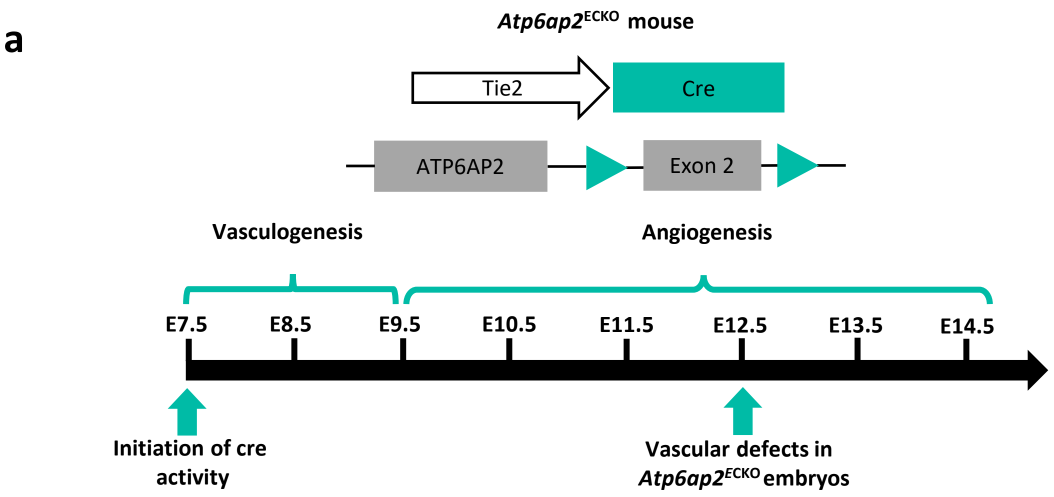


**b** **Number of P21 mice obtained from *Tie2*-Cre X *Atp6ap2* $X^f/X^f$ matings:**

| Genotype | *Atp6ap2* $X^f/Y$ | *Atp6ap2* $X^f/Y$;*Tie2*-Cre | *Atp6ap2* $X^f/X^{WT}$ | *Atp6ap2* $X^f/X^{WT}$;*Tie2*-Cre |
|---|---|---|---|---|
| Observed mice | 23 (29%) | 0 (0%) | 33 (42%) | 23 (29%) |
| Expected mice | 19.75 (25%) | 19.75 (25%) | 19.75 (25%) | 19.75 (25%) |

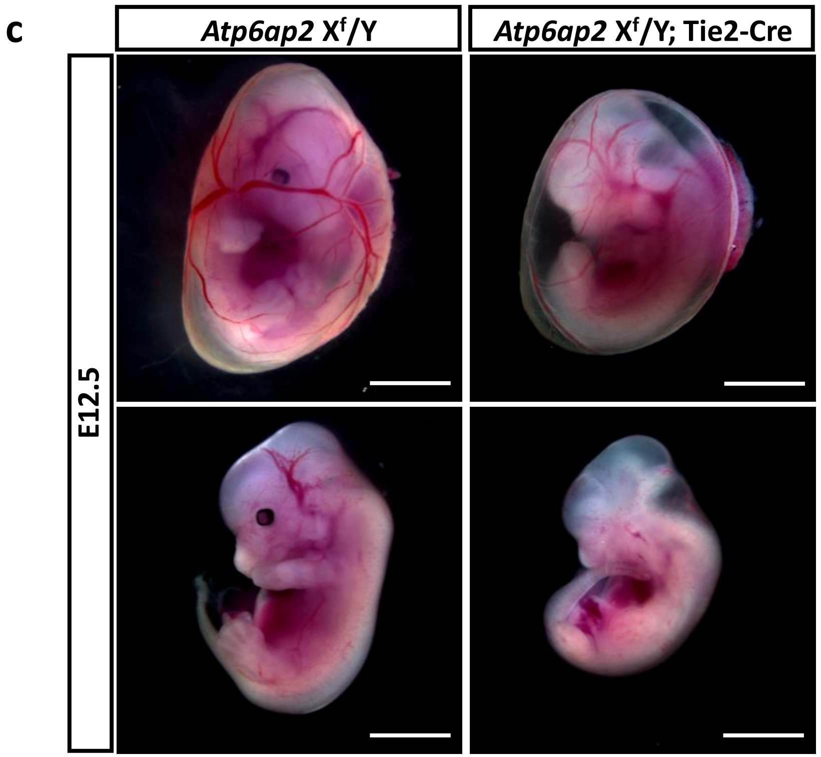

### Supplementary figure 3

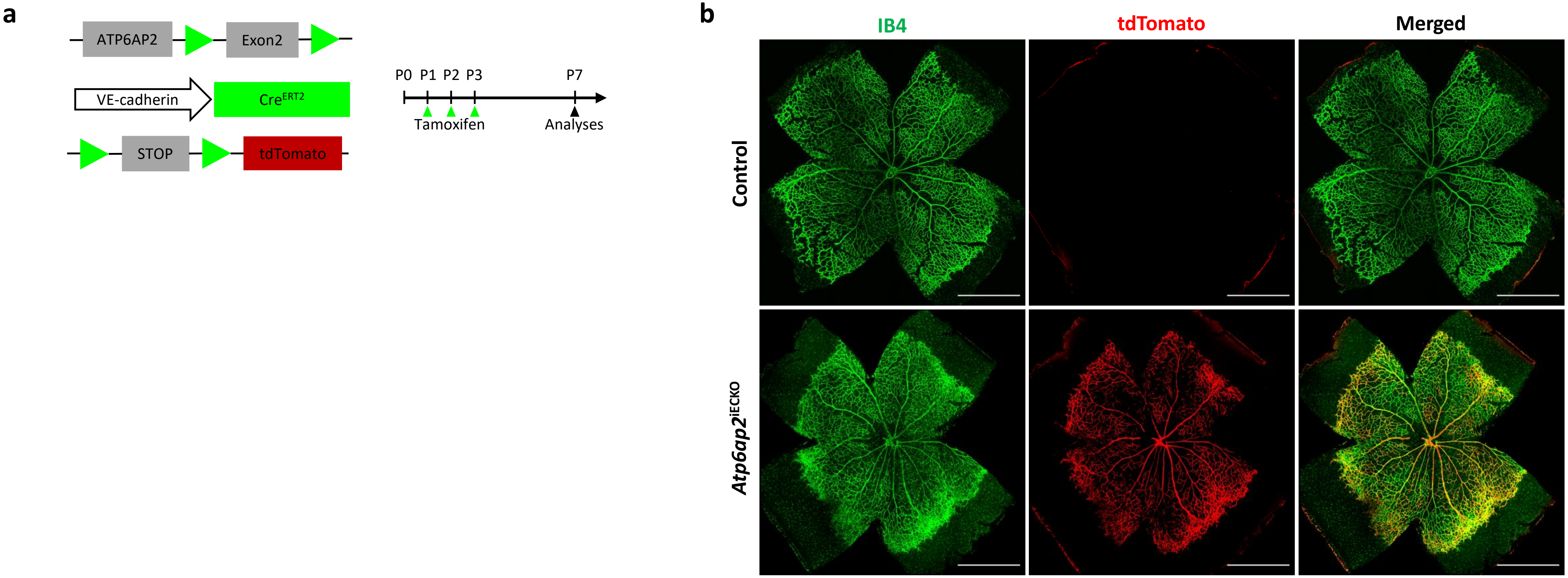

**Supplementary figure 4**

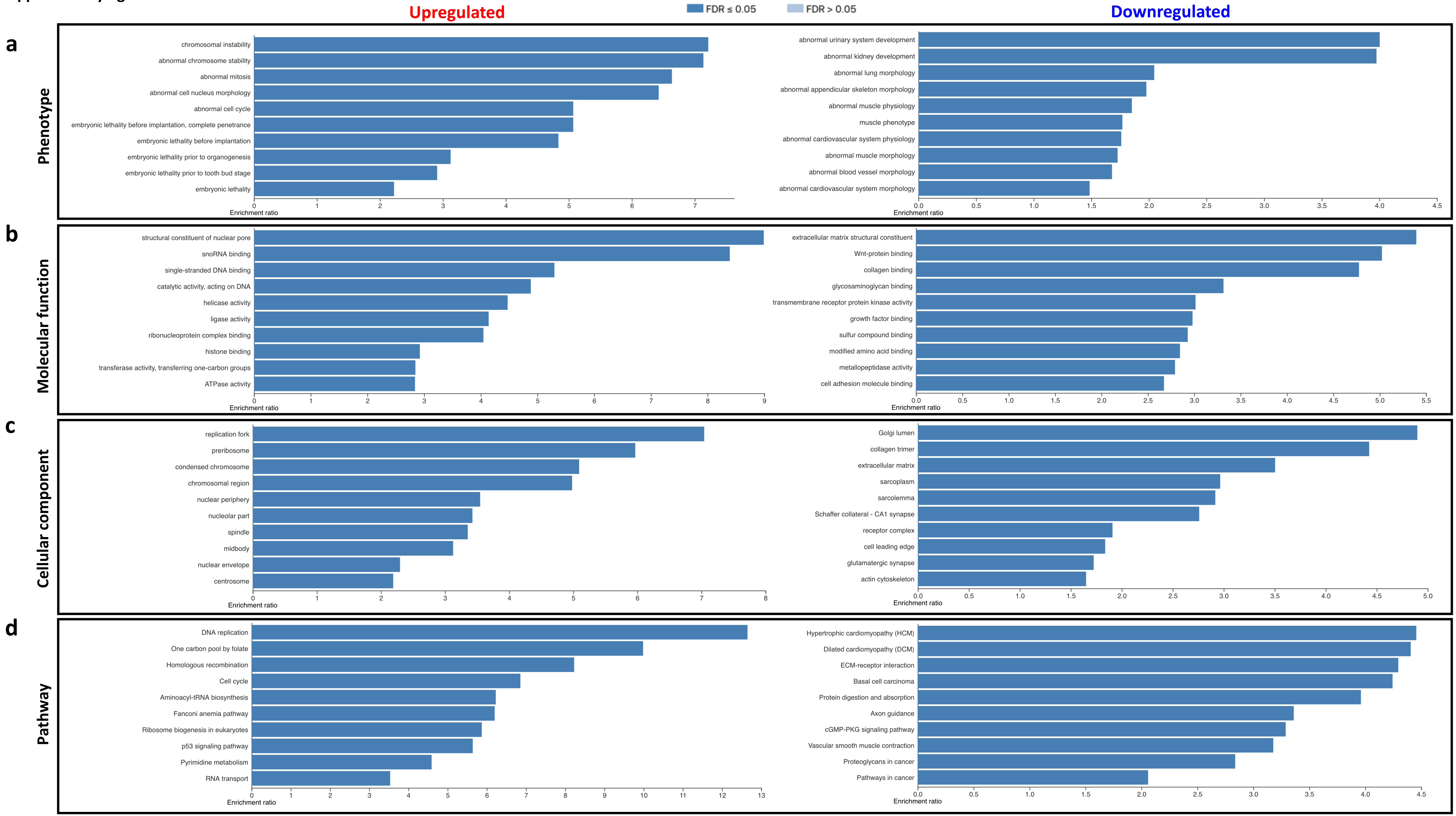

**Supplementary Table 1**
**qPCR primers**

| Gene | Forward (5' to 3') | Reverse (5' to 3') |
|---|---|---|
| *β-actin* (Mouse) | CTCTTTTCCAGCCTTCCTTCT | AGGTCTTTACGGATGTCAACG |
| *Gapdh* (Human) | TGCACCACCAACTGCTTAGC | GGCATGGACTGTGGTCATGAG |
| *Atp6ap2* (Mouse) | CCAGTTTGTTGTCTCGTCATAAGC | GCGTTCCCACCATAGAGACTG |
| *Atp6ap2* (Human) | GTGTTTTGGGGAACGAGTTTAGT | TCCTGGTATAGGCCAATTTCCA |
